# Cathepsin K maintains the number of lymphocytes *in vivo*

**DOI:** 10.1101/2020.03.04.977496

**Authors:** Renate Hausinger, Marianne Hackl, Ana Jardon-Alvarez, Miriam Kehr, Sandra Romero Marquez, Franziska Hettler, Christian Kehr, Sandra Grziwok, Christina Schreck, Christian Peschel, Rouzanna Istvanffy, Robert A.J. Oostendorp

## Abstract

In this study, we investigated the influence of the loss of Cathepsin K (*Ctsk)* gene on the hematopoietic system *in vitro* and *in vivo*. We found that cultures with Lineage^-^ SCA1^+^ KIT^+^ (LSK) cells on *Ctsk* deficient stromal cells display reduced colony formation and proliferation, with increased differentiation, giving rise to repopulating cells with reduced ability to repopulate the donor LSK and T cell compartments in the bone marrow. Subsequent *in vivo* experiments showed impairment of lymphocyte numbers, but, gross effects on early hematopoiesis or myelopoiesis were not found. Most consistently in *in vivo* experimental settings, we found a significant reduction of (donor) T cell numbers in the bone marrow. Lymphocyte deregulation is also found in transplantation experiments, which revealed that *Ctsk* is required for optimal regeneration not only of T cells, but also of B cells. Interestingly, cell non-autonomous *Ctsk* regulates both B- and T cell numbers, but T cell numbers in the bone marrow require an additional autonomous *Ctsk*-dependent process. Thus, we show that *Ctsk* is required for the maintenance of hematopoietic stem cells *in vitro*, but *in vivo, Ctsk* deficiency most strongly affects lymphocyte homeostasis, particularly of T cells in the bone marrow.

## Introduction

Secreted proteases regulate hematopoiesis through cleavage of microenvironmental factors such as critical cytokines (KITL/SCF, TGFβ) and chemokines (CXCL12), as well as matrix components and cell surface molecules required for cell-cell interactions (1). As such, proteases are instrumental in releasing hematopoietic stem cells (HSCs) from their environment, as in for instance HSC mobilization induced by granulocyte colony-stimulating factor treatment (2). Proteases include cell surface-bound dipeptidyl peptidases (including CD26 and CD143), metalloproteases (such as MMP9 and ADAMs), and cathepsins (3). The lysosomal cysteine protease cathepsin K (gene *Ctsk*, protein: CTSK) is mainly secreted by osteoclasts and cleaves collagens, osteonectin and fibrinogen, thus playing a crucial role in bone resorption (4). Deregulation of CTSK may result in osteoporosis by overactivation such as in osteoclast-mediated bone destruction in bone marrow tumors as well as in postmenopausal osteoporosis. Inactivation of CTSK by mutations in the *Ctsk* gene are further associated with increased bone density (osteopetrosis) in pycnodysostosis, an inheritable condition marked by skeletal abnormalities (5). *Ctsk* is not only expressed by osteoclasts, but it is also found in other cell types such as bone marrow (BM) stromal cells.

The role of *Ctsk* expression in BM stromal cells is poorly investigated. We have previously described that *Ctsk* is preferentially expressed by stromal cell lines which maintain repopulating HSC activity *in vitro* (6). Deletion of other stromal factors expressed by these cell lines, such as *Sfrp1* (7) or *Wnt5a* (8), results in an increased fraction of HSCs showing cell cycle progression with loss of HSC self-renewal (9). The physiologic role of CTSK in stromal cells of the hematopoietic niche has, so far, not been explored. Considering that *Ctsk* is overrepresented in HSC-maintaining stromal cell lines, we hypothesized that loss of *Ctsk* may affect HSCs and their repopulating activity in culture. To study this hypothesis, we investigate the effects of knockdown of *Ctsk* in stromal cells in co-cultures with HSCs. In addition, we explore the physiological role of *Ctsk* in WT and *Ctsk*^-/-^ mice under steady state conditions and in transplantation experiments.

## Materials and Methods

### Mice

*Ctsk* knockout mice (*Ctsk*^-/-^) (10) were bred for at least six generations on C57BL/6.J (B6) background. Age- and sex-matched B6 (CD45.2) control mice were obtained from Harlan Laboratories. In extrinsic transplantations, C57BL/6.Pep3b.Ptpcr (CD45.1) mice obtained from Taconic served as donors and recipients in intrinsic transplantations. 129S2/SvPasCrl (129; CD45.2) were obtained from Charles River Laboratories. *In vitro* experiments were conducted with (129xCD45.1)F1 (129xCD45.1) mice.

Animals were kept according to the Federation of Laboratory Animal Science Associations and institutional guidelines. All animal experiments were approved by the Government of Upper Bavaria.

### Transplantation assay

Transplantation experiments for repopulating capacity were performed as previously described (7, 8). All co-cultures were set-up with Lineage^-^ (Lin^-^) cells from 129xCD45.1 mice and transplanted into 129B6 (CD45.2) recipients. For transplantation of primary cells, 2.5×10^5^ freshly isolated BM cells were injected into lethally irradiated recipients. In extrinsic transplantations, CD45.1 WT cells were transplanted into either CD45.2 WT or *Ctsk*^-/-^ recipients. In intrinsic transplantation settings, CD45.2 WT or *Ctsk*^-/-^ BM cells were transplanted into WT CD45.1 recipients.

During the first five weeks after transplantation, recipient mice received antibiotic-treatment. Sixteen weeks after transplantation, mice were sacrificed and Peripheral Blood (PB), BM, Spleen (SPL), and Thymus (THY) were analyzed for engraftment of donor cells using the congenic system (CD45.1/CD45.2). Positive engraftment was defined as >1% positive myeloid and >1% lymphoid donor engraftment.

### Flow cytometry analysis and cell sorting

Staining of cell surface antigens was performed with antibodies from Invitrogen-ThermoFisher except for PE-Cy5.5-streptavidin conjugate, which was purchased from eBioscience-ThermoFisher. The cell suspension was incubated in HF2+ buffer (Hank’s balanced salt solution with 2% FCS and 10 µM HEPES buffer) for 15 min on ice in the dark. Hematopoietic subpopulations were separated as described previously (7, 8). In some experiments, a combination of both CD4 and CD8a antibodies was used to stain for T cells. The results of these were indistinguishable from those where we used only CD3ε. Thus, we have combined the results of all experiments and report these as CD3ε+ throughout the manuscript. EPICS XL (Beckman Coulter) or CyAn ADP Lx P8 (Coulter-Cytomation) were used for FACS analysis; data were analyzed using FlowJo software (Tree Star). Cell sorting was done with a MoFlo High Speed cell sorter (Beckmann Coulter).

### Knock down of *Ctsk* in UG26-1B6 stromal cells and cocultures with HSCs

UG26-1B6 stromal cells were cultured on 0,1% gelatin coated wells as described (11). As previously published (7, 12), we used established stable *Ctsk* gene knockdowns in stromal cells. In brief, sh*RNA*^mir^ contructs on the pLKO.1 backbone for *Ctsk* gene knockdown (Open Biosystems - ThermoFisher) were transfected into Phoenix E packaging cell line. Collected virus particle were used for the infection of UG26-1B6 cells followed by selection with 5 µg/ml puromycin. As a control, empty vector pLKO.1 was used.

For cocultures, UG26-1B6 derivatives sh*Ctsk2 and -4*, and pLKO.1 were grown in 6 well plates to 90% confluence and irradiated with 30 Gy. 1.0×10^4^ Lin^-^ BM cells were plated on stromal cells and cultured in long–term culture medium (M5300; Stem Cell Technologies) supplemented with hydrocortisone in a 1:1000 ratio, Penicillin/Streptomycin and GlutaMax (200mM, GIBCO-ThermoFisher) in a 1:100 ratio for two to four weeks. Each week half of the supernatant was replaced with fresh medium. After coculture, cells were either used for colony forming unit and FACS analysis or for *in vivo* repopulation assay.

### Hematopoietic Colony Assays

Colony forming assay was performed with growth factor-supplemented methylcellulose (Metho Cult GF 3434, STEMCELL Technologies). 2.5×10^4^ total BM cells, 1×10^4^ Lin^-^ BM cells or fractions of cell cultures were seeded into methylcellulose on two 3.5 cm dishes. After ten days of culture at 37°C, 5 % CO_2_ and ≤95% humidity, the colonies formed were counted under the microscope and analyzed using flow cytometry.

### Single cell culture

Serum-free conditioned medium (CM) was essentially generated as described (8, 12, 13) by incubating serum-free medium (BIT 9500, STEMCELL Technologies) for three days on irradiated (30 Gy) and confluent pLKO.1 and sh*Ctsk* stromal cells.

Single cell cultures were performed as previously published (8, 12, 13). Briefly, HSC-enriched CD34^-^ CD48^-^ CD150^+^ Lin^-^ SCA1^+^ KIT^+^ (CD34^-^ SLAM) cells were sorted from the lineage-depleted BM cells of 8 weeks old B6 control mice and deposited as single cells in round-bottomed 96-well plates. Each well was filled with 100µl of 0,22µm filtered CM, supplemented with 100ng/ml mSCF and 20ng/ml IL-11 (both from R&D Systems). The clone size was counted each day under the light microscope. After five days, cells were harvested and analyzed by flow cytometry.

### Statistical methods

Significance levels were determined with unpaired Students t-test when data followed a Gaussian distribution. When data were not normally distributed, we used the Mann-Whitney test analyzed by GraphPad Prism 7.

## Results

### Reduction of stromal Cathepsin K impairs proliferation and clonogenic capacity of HSCs *in vitro*

To study how the absence of stromal *Ctsk* affects hematopoiesis *in vitro*, we generated *Ctsk*-knockdowns in the HSC supportive stromal cell line UG26-1B6 with sh*RNA*^mir^ (sh*Ctsk2* and sh*Ctsk4*). As control cells, we transduced stromal cells with pLKO.1 empty vector backbone. The *Ctsk* mRNA expression was reduced to 70% in sh*Ctsk*4 and till 90% in sh*Ctsk*2 as compared with pLKO.1 controls (Figure 1A). Also, CTSK protein content was reduced by 60% in sh*Ctsk*2 stromal cells (Figure 1B).

**Figure 1.**
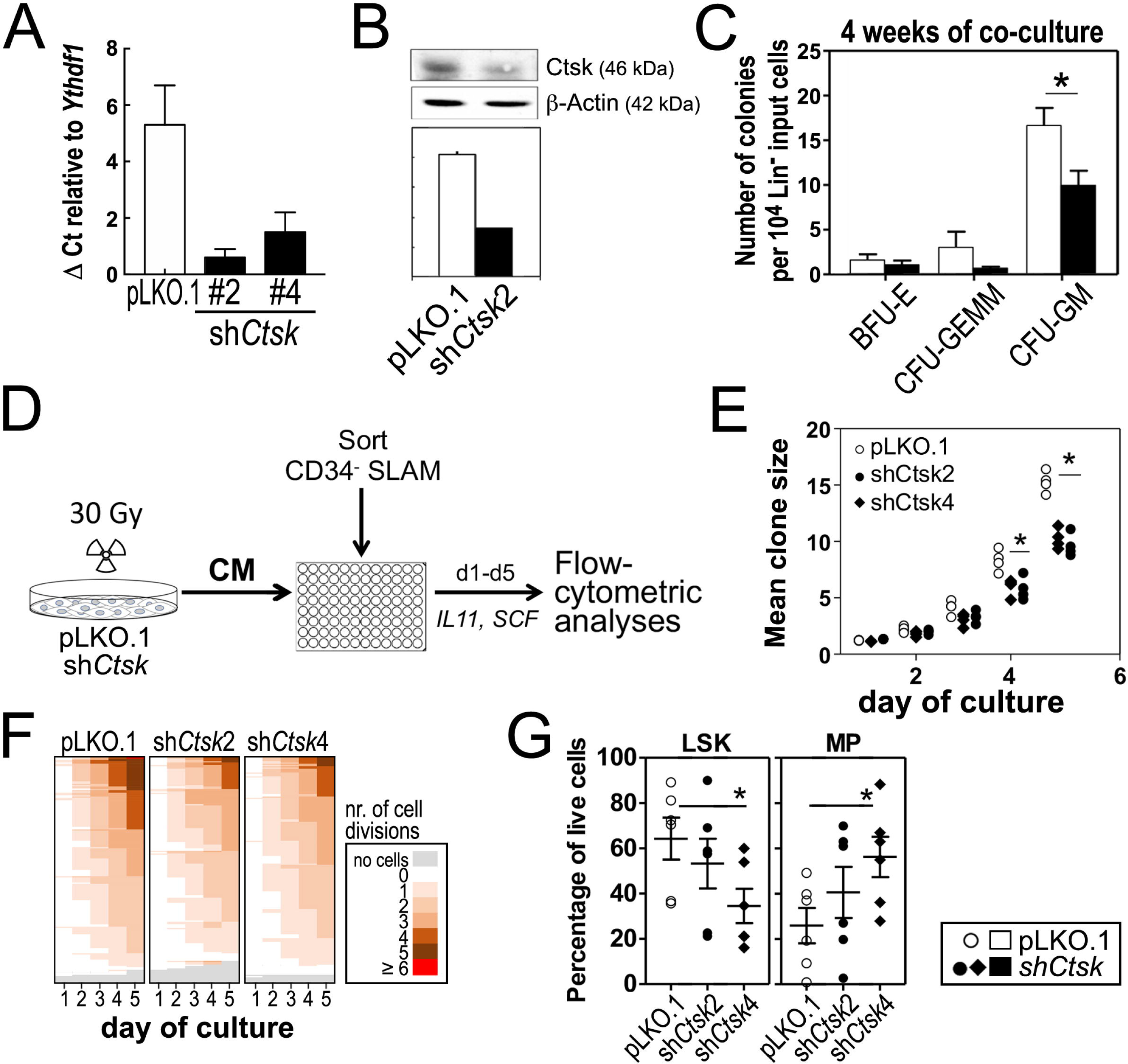
*In vitro* experiments with UG26-1B6 -pLKO.1 and -sh*Ctsk* knockdown stromal cells. **(A)** Relative mRNA level, measured by real-time PCR (n=3) using *Ythdf1* as housekeeping gene, comparing the relative expression of *Ctsk* in the knockdown clone of the UG26-1B6 (sh*Ctsk2 and shCtsk4*) cells and in the pLKO.1 empty vector control. *ShCtsk2* was used in the follow-up experiments, unless otherwise indicated **(B)** Western Blot showing the relative CTSK protein content to ß- Actin in UG26-1B6-pLKO.1 and sh*Ctsk2* cells (n=3, quantified by ImageJ). **(C)** Co-culture of 10.000 Lin^-^ cells on irradiated *shCtsk* and pLKO.1 stromal cells in 10 cm^2^ round dishes for 4 weeks (n=6). After culture, cells were seeded in methylcellulose, after ten days colonies were counted under the microscope. **(D)** Experimental design of single cell experiments: Conditioned medium was generated as described in the Materials and methods section. Single CD34^-^ SLAM cells were sorted in 96-well plates with pLKO.1 and *shCtsk2* and *shCtsk4* CM supplemented with SCF and IL11 and every 24h microscopically evaluated for cell number. After five days (d1-d5), the clones were harvested and analyzed using flow cytometry (n=2). **(E)** Mean clone size of CD34^-^ SLAM cells cultured with pLKO.1 and *shCtsk* CM. Shown are the results of four independent experiments. **(F)** Heat map showing proliferation frequency of single clones colored by number of divisions. **(G)** Content of LSK and MP cells per cultured plate. Cells from all wells were pooled and stained for lineage markers (CD45R[B220], CD4, CD8a, CD11b, and Gr1), SCA1 and KIT. Shown are the results of two independent single cell experiments, each with three plates (CD34^-^ SLAM cells from separate mice) per condition. *: p<0,05 using Mann-Whitney U test. Open symbols: cultures with control (pLKO) stroma or CM, closed symbols: cultures with sh*Ctsk* stroma or CM.

To determine whether reduction of *Ctsk* mRNA affects production of colony-forming cells, we set up co-cultures of WT Lin^-^ BM-cells with pLKO.1 and sh*Ctsk2* stromal cells. These experiments showed, that four weeks after co-culture the number of granulocytic colony-forming cells (CFU-GEMM and CFU-GM) was reduced, when stromal *Ctsk* was down-regulated (Figure 1C). To determine whether the decreased production of granulocytic progenitors directly affected HSC-enriched CD34^-^ SLAM cells, we performed stroma-free single cell cultures supported by stromal conditioned medium (CM) from pLKO.1 and sh*Ctsk* stromal cells supplemented with SCF (KIT-Ligand) and IL-11 (Figure 1D). In these cultures, proliferation rates and clone sizes of CD34^-^ SLAM cells cultured in either *shCtsk2 or shCtsk4* CM were significantly reduced compared with cells cultured in CM from pLKO.1 control (Figure 1E, 1F). Importantly, the frequency of LSK cells was reduced in both sh*Ctsk* CMs (20%) compared with pLKO.1 CM (45%), while myeloid progenitors (MPs) frequency was increased 1.2-to 2.3-fold after culturing in CM from both sh*Ctsks* (Figure 1G). Despite a slightly higher knockdown of the *Ctsk* gene in *shCtsk2* compared to *shCtsk4* (Figure 1A), both stromal cells affected hematopoiesis in a similar manner in cultures. Our findings suggest an enhanced differentiation towards more mature MPs in CM from sh*Ctsk* stromal cells with a slow loss of the more undifferentiated LSK cells.

To analyze whether the slow loss of undifferentiated cells in culture also affected cells with repopulating ability, we transplanted three-weeks co-cultures of sh*Ctsk* and pLKO.1 stromal cells and WT Lin^-^ BM cells into WT recipient mice (Figure 2A). Since in previous experiments results on sh*Ctsk2* and sh*Ctsk4* stromal cells were very similar, In these and all subsequent cultures, we only used the sh*Ctsk2* as sh*Ctsk* cells. Unexpectedly, cells from both types of culture showed similar total engraftment in the peripheral blood (PB, Figure 2B). In addition, overall donor myeloid and lymphoid engraftment in the PB was also unchanged between transplants od pLKO.1 and sh*Ctsk2* co-cultures (not shown). Similarly, donor engraftment in the BM of recipient mice was also unchanged. Interestingly, the only mature population affected after transplantation of Lin^-^ cells cultured on sh*Ctsk* stroma were donor T (CD3ε^+^) cells, which were reduced by 60% in the small T lymphocytic BM compartment of recipient mice (Figure 2C). Furthermore, despite lack of alteration in myeloid populations in the PB, regeneration of the donor HSC-enriched fraction of LSK cells was significantly reduced in the BM (Figure 2D).

**Figure 2.**
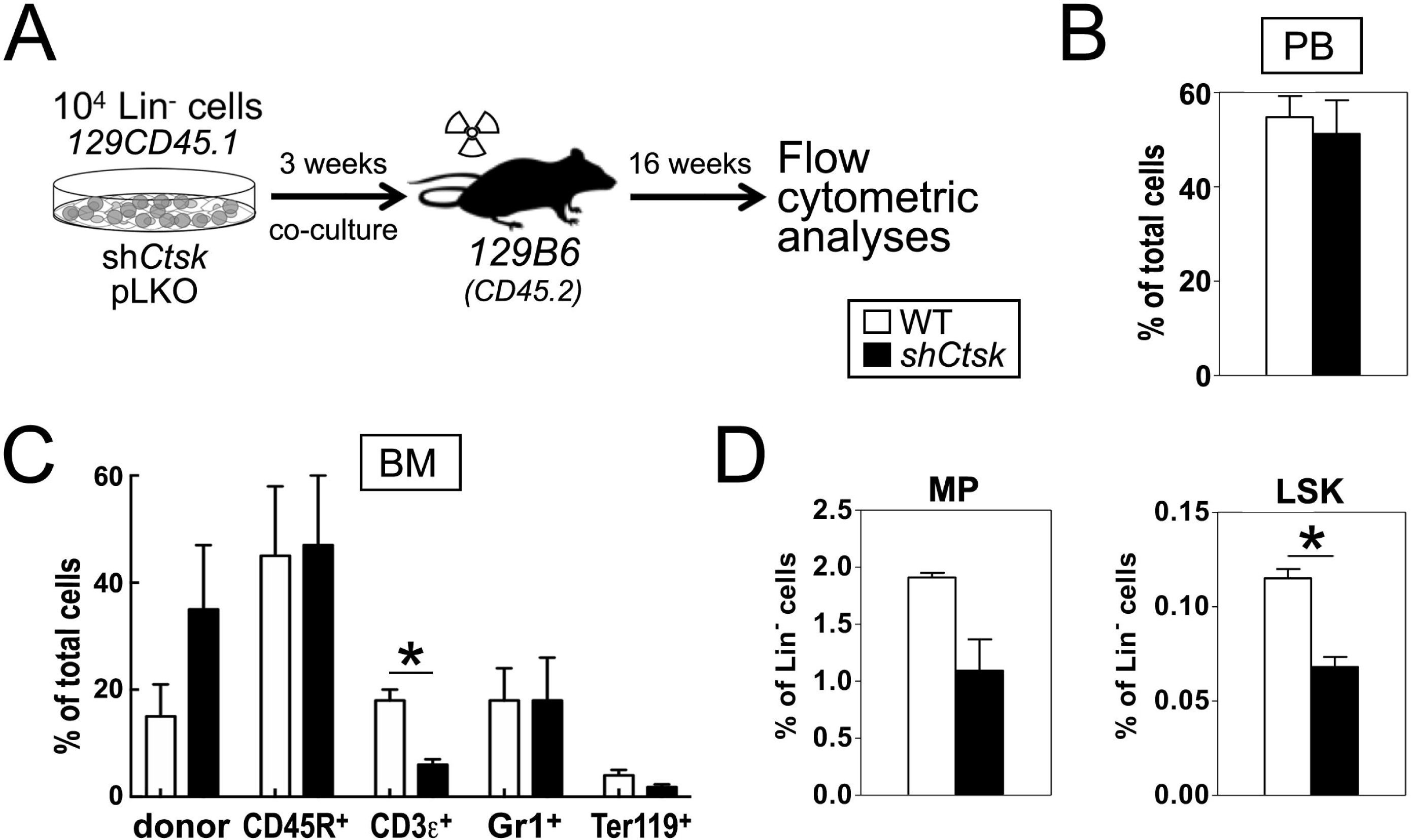
Maintenance of repopulating activity in cocultures of HSC and UG26-1B6 - pLKO.1 and -sh*Ctsk* knockdown stromal cells. **(A)** Experimental design of co-culture transplant experiments: 10.000 Lin- cells were co-cultured on irradiated stromal cells for three weeks and then transplanted into lethally irradiated mice (two independent experiments totaling UG26-1B6-pLKO: n=7; UG26-1B6-sh*Ctsk*: n=6 recipient mice) **(B)** FACS analysis of PB, 16-weeks post transplantation, showing the level of donor engraftment as percentage **(C)** FACS analysis of BM, 16-weeks post transplantation, showing percentages of donor-derived mature and **(D)** early stage hematopoietic cells. *: p<0,05 using Mann-Whitney U test. Open bars: co-cultures on control stroma; closed bars: co-cultures on sh*Ctsk* stroma.

### Loss of Cathepsin K modulates *in vivo* lymphocyte homeostasis

Since reduction of stromal *Ctsk* reduces both number and function of early hematopoietic cells *in vitro*, we wondered whether *Ctsk* deletion similarly affects hematopoiesis *in vivo* in *Ctsk* knockout (*Ctsk*^-/-^) mice. In 8-12 weeks old *Ctsk*^-/-^ mice we found that early hematopoiesis and myelopoiesis was unchanged under steady state conditions compared to WT controls (Figures 3A, B). When evaluating lymphocytic cells, we found that the fraction of peripheral CD45R^+^ (B220^+^ B) cells and also the number of the small population of thymic B cells were both significantly elevated (Figures 3C, D). Also in the thymus, the number of CD3ε^+^ thymocytes was increased by 2-fold (Figure 3D) with a reduction in the double-negative CD4^-^ CD8a^-^ T cell population (Supplementary Figure 1). In contrast, in the spleen (SPL), the numbers of B and T cells were neither increased nor reduced (Figure 3E). Interestingly, consistent with the results of transplanted cells from sh*Ctsk* co-cultures, we found a reduction in the small fraction of CD3ε^+^ (T) cells of about 60% in the BM (Figure 3F). These results suggest a deregulation in distribution of both B- and T-lymphocytes over different hematopoietic tissues in the absence of *Ctsk*.

**Figure 3.**
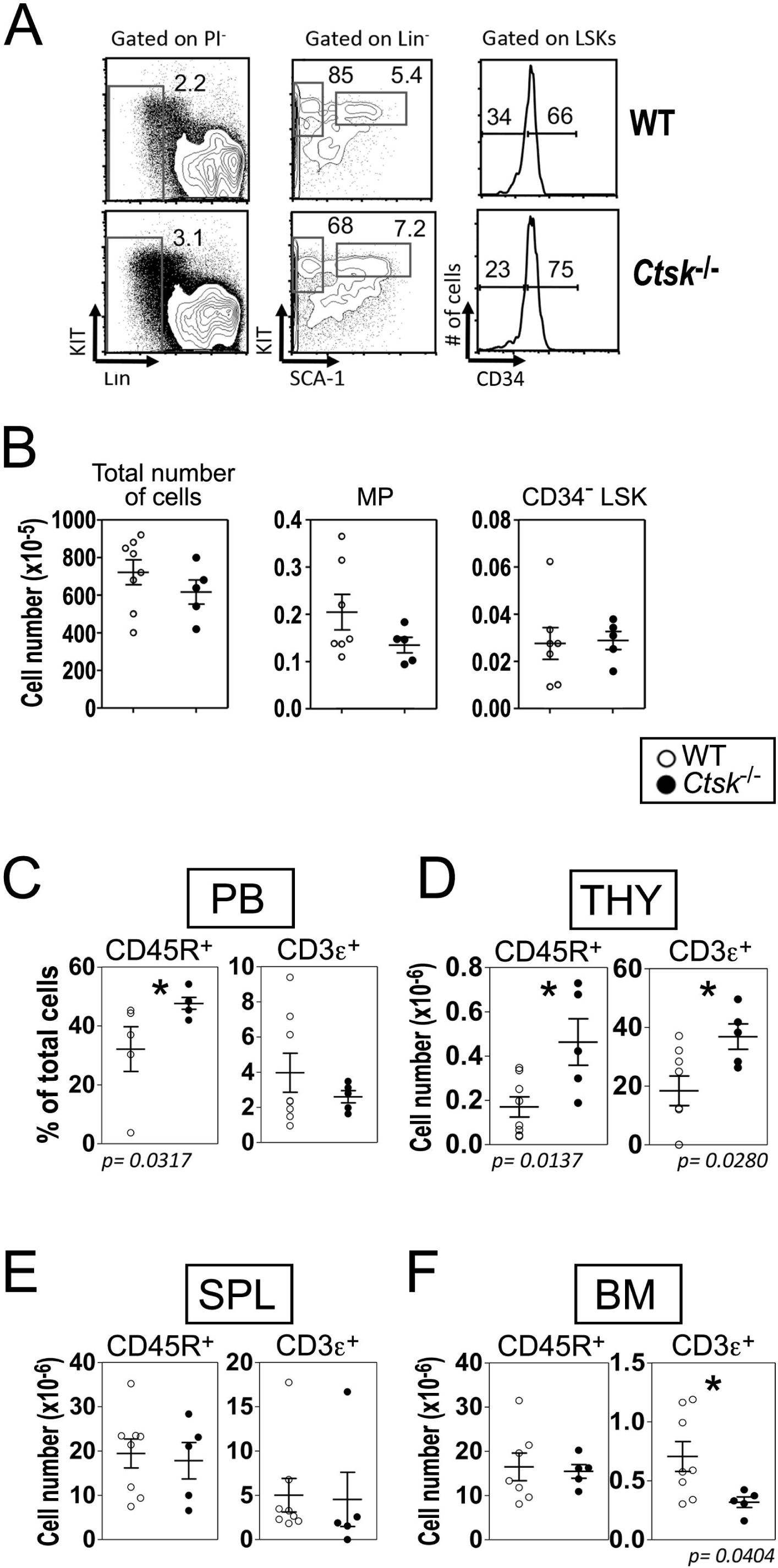
Effects of *Ctsk* loss on steady state hematopoiesis. (**A**) Representative FACS plots of the BM stem and progenitor cells from the BM of WT and *Ctsk*^-/-^ mice. (**B**) Absolute numbers of cells, myeloid progenitors and CD34^-^ LSK cells in the BM of WT and *Ctsk*^-/-^ mice. (**C**) Percentages of B- and T-lymphocytes in the PB, and (**D**) absolute numbers of B and T cells in the BM (**D**), spleen (SPL, **E)** and thymus (THY, **F**). Data represents results of two to three independent experiments. In B-E each dot represents one animal. Open symbols represent results from WT control animals, and closed symbols represent *Ctsk*^-/-^ animals. *: p<0,05 using the Mann-Whitney U test. Open symbols: WT mice (*Ctsk*^+/+^ littermates); closed symbols: *Ctsk*^-/-^ mice.

### *Ctsk* is required for regeneration of donor lymphocyte numbers

In the next series of experiments we studied whether the knockout of *Ctsk* in the microenvironment would affect the maintenance of WT HSCs in transplantation settings (7, 8). For this purpose, we transplanted WT (CD45.1) BM cells into lethally irradiated *Ctsk*^-/-^ (CD45.2) mice or their WT littermates (Figure 4A). These experiments showed that 16 weeks after transplantation, a comparable peripheral engraftment of donor cells was given (Figure 4B). In contrast to the experiments in which co-cultures on sh*Ctsk* stromal cells were evaluated (Figure 2), overall regeneration of HSC-enriched CD34^-^ LSKs and myeloid progenitors (MPs) in the BM of WT and *Ctsk*^-/-^ recipient mice appeared to be unaffected (Figures 4B-D). When evaluating lymphocytic populations, we found that in accord with unchanged peripheral engraftment, B- and T-cell populations were also unchanged in the periphery (not shown). However, we found deregulation of donor B- and T-lymphocyte numbers in several hematopoietic tissues of *Ctsk*^-/-^ recipient mice as compared with their WT littermate recipients. In particular, we found reductions of both CD45R^+^ B and CD3ε T cells in the BM and Thymus (THY) (Figure 4E, F). In the THY, the number of donor-derived CD3ε^+^ cell repopulation was strongly decreased by almost 70% in *Ctsk*^-/-^ recipient mice as compared with their WT littermates (Figure 4F), suggesting a role for *Ctsk* in regeneration of the BM and THY microenvironments *in vivo*. Furthermore, despite reductions in BM and THY, the number of CD45R^+^ B cells was increased in the spleens of *Ctsk*^-/-^ recipient mice (SPL) (Figure 4G; Supplementary Figure 2). These experiments show that *Ctsk* is required for optimal repopulation and maintenance of donor lymphocyte numbers.

**Figure 4.**
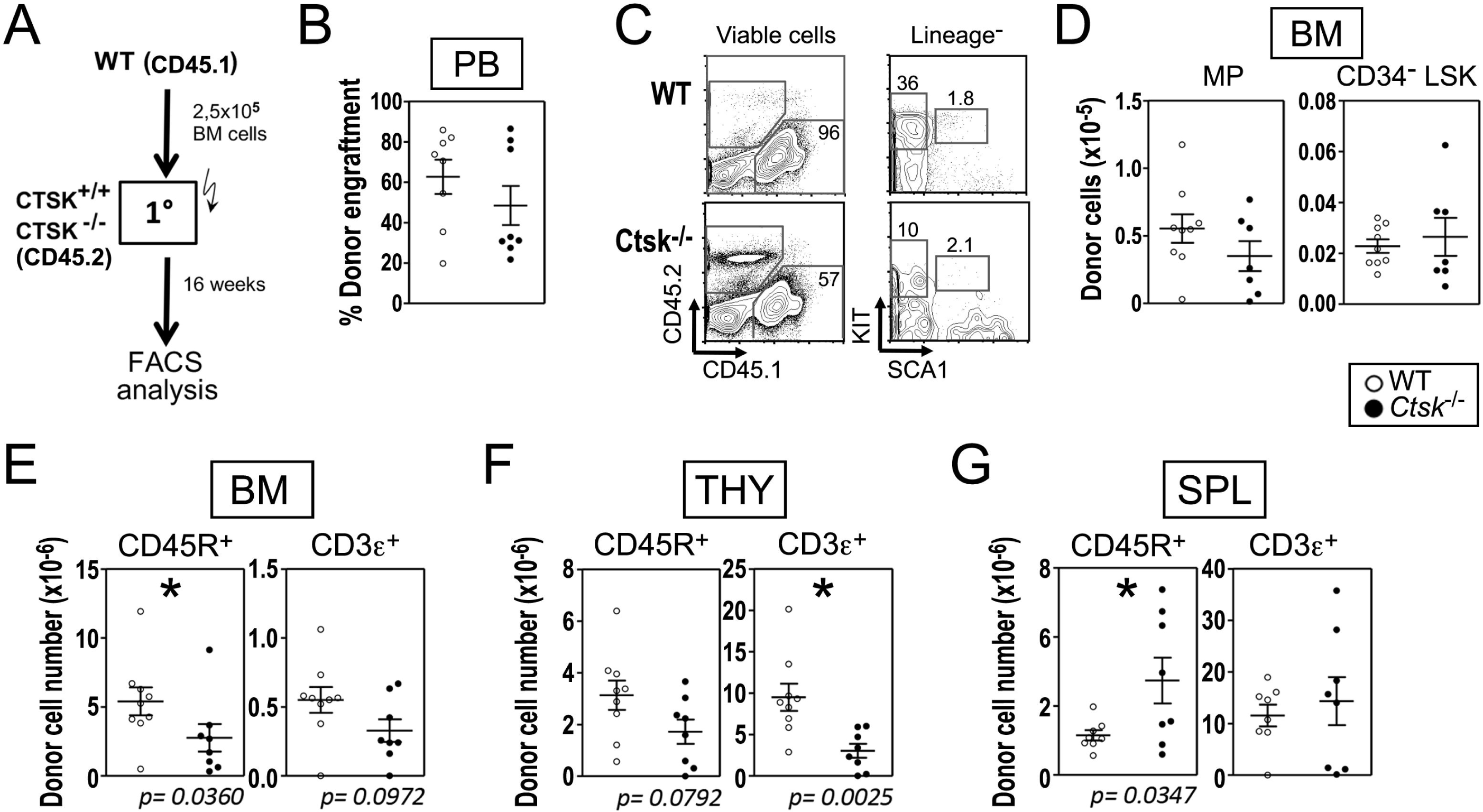
Role of extrinsic *Ctsk* for regeneration of WT HSCs. (**A**) Experimental design. WT Lin^-^ cells from congenic CD45.1 mice were transplanted into CD45.2 *Ctsk*^-/-^ mice and their WT littermates. Donor cells were analyzed 16 weeks post transplantation. (**B**) Percentage of donor engraftment in the PB. (**C**) Representative FACS plots of the BM stem and progenitor cells. (**D**) Absolute numbers of MPs and CD34^-^ LSKs in the BM from the extrinsic transplant recipients. (**E**) Absolute numbers of B- and T- lymphocytes in the BM, (**F**) SPL, and (**G**) THY. Results of two or three independent experiments are shown. In B, D-G, each dot plot represents one animal. Open symbols represent results from WT control animals, and closed symbols represent *Ctsk*^-/-^ recipients. *: p<0.05 statistically significant using Mann-Whitney U-test. Open symbols: WT recipient mice; closed symbols: *Ctsk*^-/-^ recipient mice.

To determine whether the lymphocyte deregulation was caused by cell-extrinsic or cell-intrinsic loss of *Ctsk*, we transplanted HSCs from Ctsk^-/-^ mice into lethally irradiated WT recipient mice (Figure 5A). Interestingly, 16 weeks after transplantation, we found that *Ctsk*-deficient BM cells poorly repopulated the periphery of recipient mice (Figure 5B). But, this reduction in donor cells was not due to impaired hematopoiesis, since engraftment of early donor hematopoietic compartment was unaltered in the BM of recipient mice transplanted with *Ctsk*^-/-^ cells, suggesting that myelopoiesis was unaffected (Figure 5B, C). With regard to lymphocytic compartments, we found, in accord to extrinsic transplantation results of WT HSC into a *Ctsk*^-/-^ environment (Figure 4F), a reduction of the small T lymphocyte fraction in the BM (Figure 5D, Supplementary Figure 3). But, other hematopoietic organs, like SPL (Figure 5E) and THY (data not shown) showed no significant changes in the number of donor-engrafted lymphocytes.

**Figure 5.**
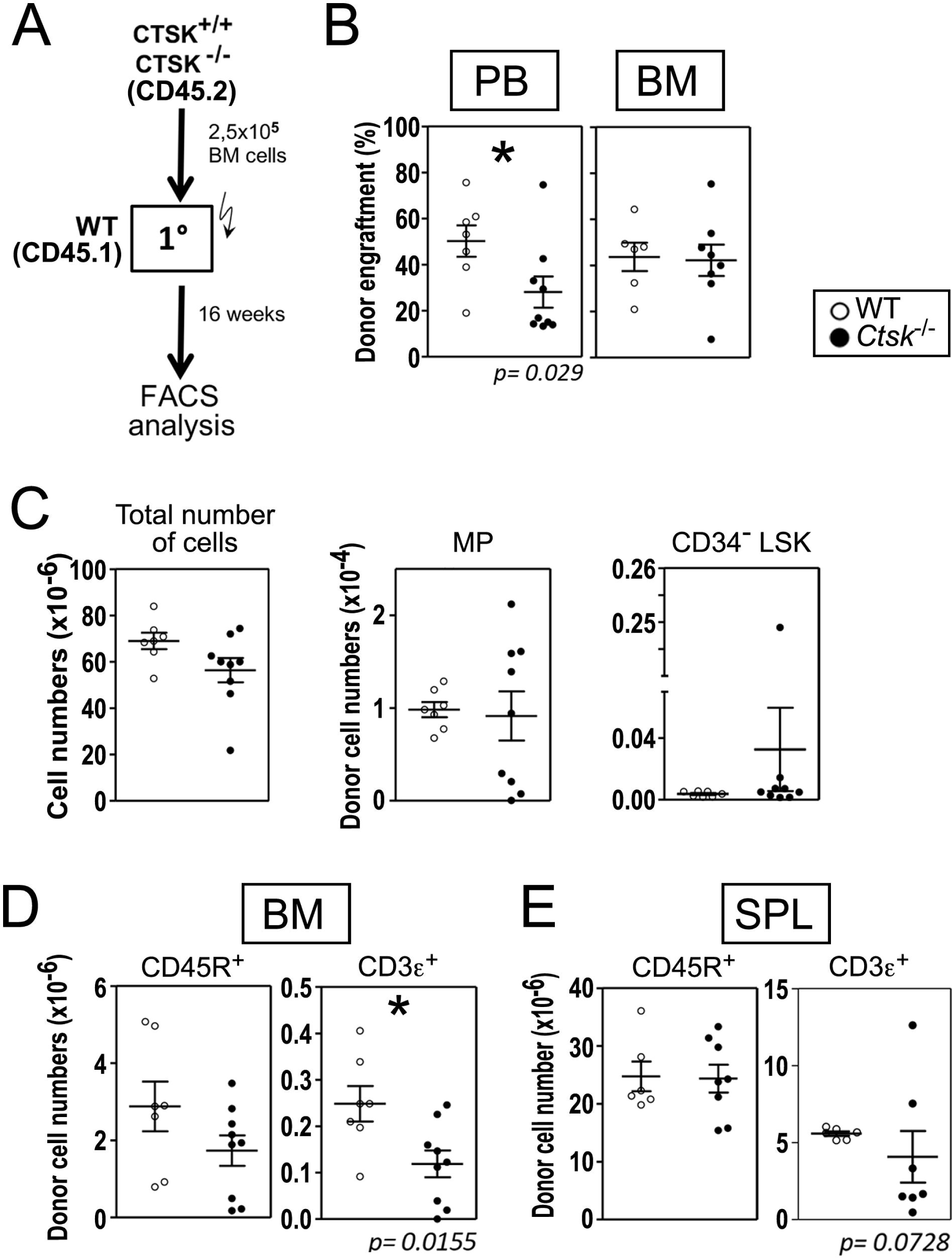
Repopulation capacity of *Ctsk*^*-/-*^ HSCs. (**A**) Experimental design. CD45.2 BM cells from *Ctsk*^-/-^ and WT control littermates were transplanted into CD45.1 WT recipient mice and analyzed 16 weeks post transplantation. (**B**) Percentage of donor engraftment in PB, and BM. (**C**) Absolute numbers of donor cells, MPs, and CD34^-^ LSKs in the BM of recipient mice. Absolute numbers of donor B and T cells in the BM (**D**) and SPL (**E**). The results of two independent experiments are shown. In B-E, each dot represents one animal. Open symbols represent results from WT donor animals, and closed symbols represent *Ctsk*^-/-^ donors. *: p<0.05 statistically significant difference between control and *Ctsk*^-/-^ donor cells in WT recipients using Mann-Whitney U-test. Open symbols: WT donor mice; closed symbols: *Ctsk*^-/-^ donor mice.

## Discussion

We explore the role of stromal *Ctsk* in the maintenance of HSCs *in vitro* and its importance for the regulation of steady state hematopoiesis and the number of regenerating lymphocytes in transplantation experiments *in vivo*. Our experiments show that *Ctsk* expression is required for maintenance and proliferation of HSCs *in vitro* and was particularly important for the maintenance of cells regenerating the small fraction of regulatory BM T cells. Despite these clear indications of the importance of *Ctsk* for maintaining HSCs in culture, we found that *Ctsk* loss in vivo did not significantly affect HSC maintenance and myelopoiesis *in vivo*. Remarkably, our experiments indicate that *Ctsk* regulates the regeneration of B- and T-cell numbers in multiple organs. Importantly, while B cell repopulation mainly depends on cell-extrinsic *Ctsk* expression, the CD3ε^+^-cell compartment in the BM is affected both by cell autonomous and non-automonous *Ctsk* deficiency.

We found that knockdown of *Ctsk* in stromal cells reduces their ability to maintain the number and function of HSC in culture, resulting in reduced production of granulocytic progenitors, lower rates of HSC proliferation as well as the maintenance of their ability to repopulate MP and LSK compartments. Considering that CTSK is a cysteine protease, this result suggests that degradation of CTSK substrates is required for maintaining HSC in culture. One CTSK substrate is collagen. We previously found that collagen-mediated activation is upregulated in HSCs exposed to stromal factors (13). More importantly, together with neural growth factor, collagen could replace stromal factors to ensure optimal survival of HSCs in culture (13). However, here we found no effect of *Ctsk* depletion on the survival-promoting activity of stromal CM *in vitro*, suggesting that the specific HSC survival promoting collagen is not affected by CTSK, or that collagen may be substituted by other stromal factors to maintain HSC *in vitro*. Other CTSK substrates were shown to be SCF, osteopontin, and the chemokine CXCL12 (3), which all have documented activity on HSCs and which are required for HSC maintenance. However, in previous studies revealing the transcriptome of the HSC-supportive UG26-1B6 cells, we did not detect expression of these three factors (12, 13), suggesting that in co-cultures with UG26-1B6, these three factors may not be involved. Thus, it will now be interesting to figure out the relevant CTSK substrates for HSC maintenance on stromal cells in vitro by secretome analysis.

In contrast to the mild effects of *Ctsk* deletion on *in vivo* myelopoiesis, we here uncovered a consistent imbalance in the numbers and distribution of T-cells in four different experimental settings in different hematopoietic tissues. Importantly, we describe deregulation under both steady-state and transplantation conditions. We found both up- and down-regulation of T cell numbers under different conditions. Most consistently, we observed a reduction of the number of T cells in the BM in all experimental models explored. The picture is not so clear in other tissues. For instance, under steady state conditions, CD3ε^+^ cell numbers in the thymus increase significantly with a particular increase in CD4^+^ cells. The mechanism here is unclear but could be caused by decreased T-cell selection or reduced cell death. In contrast, under stress conditions, such as regeneration of donor T cell numbers in *Ctsk*^-/-^ mice transplanted with HSCs, the number of T-cells is reduced in the thymus. This observation suggests a differential requirement for *Ctsk* expression under steady state conditions and situation of regenerative stress. Although the upregulation of thymocytes under steady state conditions remains enigmatic, reduced regeneration of T cells could have several causes. For instance, successful transplantation requires myeloablation, which in our case, was achieved by lethal irradiation. *Ctsk* might play a role in the regeneration of the thymic microenvironment, which is impaired in *Ctsk*^-/-^ mice after irradiation. Furthermore, as noted, Ctsk is not only important in cell non-autonomous regulation, but also in autonomous processes. In this respect, it has been shown that cathepsins play a role in endosomal processing required for antigen presentation, which have been described in mice deficient in cathepsin B (14). The absence of Ctsk could, in analogy, lead to dysfunctional presentation of antigens in the context of MHC Class II or CD1d molecules and defective Th2 lymphocyte responses (14). In this context, the report that FoxP3^+^ regulatory T cells from *Ctsk*^-/-^ mice were much more potent in suppressing T effector cells (15) could also be considered.

The most consistent finding of our study is the reduction of CD3ε^+^ cells in the BM, which is found under steady state conditions as well as in transplantation of cultured cells and in both transplantation of WT into *Ctsk*^-/-^ recipients and vice versa. The regulatory role of the small marrow population of *Ctsk*-sensitive CD3ε^+^ cells is unclear at the moment. This small cell population in the BM might play an important role in the regulation of hematopoiesis. Indeed, it has been reported that the population of FoxP3^+^ Tregs accounting for 0.15% of total BM, which is slightly less than the number of HSC (16), preserve HSC quiescence and stimulate engraftment (16). Together with the abovementioned increased potency of Tregs from *Ctsk*^-/-^ mice (15), it would be of interest to establish what mechanisms cause the reduction in BM T-cells, and particularly FoxP3^+^ Tregs, and how these cells localize and regulate myelo- and lymphopoiesis in *Ctsk*-deficiency.

In addition to imbalances in T-cells, we also find in some settings an imbalance in the number of CD45R^+^ B-cells. Under steady state conditions these cells increase in the periphery, and they also increase of 2.7-fold in the thymus. As is the case of the T cell compartment in the BM, thymic B cells represent only a minor population in the thymus. Like T cells in the thymus, thymic B cells are increased in steady state, but they show decreased repopulation in the thymus after transplantation. Whereas, the reduction of thymic B cells could be caused by impaired regeneration of the thymic microenvironment in the absence of *Ctsk*, the reason for the increase of the thymic B cells under steady state conditions remains to be established. This small B cell population has been shown to play a pivotal role in negative selection of T cells (17) and it promotes development of regulatory T cells in the thymus (18). Their development depends on both hematopoietic-intrinsic and thymic microenvironment-intrinsic regulatory mechanisms (19). Our results suggest that either their development or the homing of B cell precursors into the thymus from the periphery is influenced by CTSK. The latter possibility appears likely, since in the experiments where we transplanted WT HSC into *Ctsk*^-/-^ mice, the decline in CD45R^+^ cells in the BM and thymus was associated with an increase in the spleen, suggesting that the total number of B cells in the *Ctsk*^-/-^ recipients was not altered.

In conclusion, *Ctsk* is expressed by stromal cells of the hematopoietic niche. We here describe that *Ctsk* plays an important role in the maintenance of CD34^-^ SLAM cells and their myeloid differentiation *in vitro*. In addition, stromal *Ctsk* deficiency decreases number and function of cells able to regenerate the LSK compartment after transplantation *in vivo*. Despite these effects of Ctsk deficiency *in vitro* defects in early hematopoiesis and myelopoiesis are only mild in *Ctsk*^-/-^ mice, suggesting a redundancy for the requirement of CTSK *in vivo* which is not observed *in vitro*. The main effects of *Ctsk* deletion *in vivo* reveals a recurrent reduction in T lymphocytes in the BM as well as fluctuations of both B- and T-cell numbers compared to WT animals or recipients in different organs at steady state and in transplantation experiments. Importantly, in transplantation experiments, both autonomous and cell non-autonomous regulation by *Ctsk* deficiency can be observed. Thus, our study shows that intact activity of CTSK is required for *in vitro* HSC maintenance in stromal co-cultures. In contrast, *in vivo*, myelopoiesis is only mildly affected by *Ctsk* deficiency, but CTSK maintains B and T cell numbers in different hematopoietic organs, where in- or decreases in cell numbers most probably depend on the effects of stress on the microenvironment. The most consistently observed role of CTSK is to non-redundantly regulate T cell numbers in the BM.

## Supporting information

Hausinger CTSK Supplemental Files

## Acknowledgements

We thank Prof. Paul Saftig (Unit for Molecular Cell Biology and Transgenic Research, Department of Biochemistry, University of Kiel, Kiel, Germany) and Prof. Ulrich A. Maus (Division of Experimental Pneumology, Institute of Immunology, Medical University Hannover, Hannover, Germany) for supplying the *Ctsk*^-/-^ for this study. We further thank Sandra Grziwok, Charlotta Pagel and Alina Wagner for expert technical assistance. We further thank Lynette Henkel and Dr. Matthias Schiemann (CyTUM-MIH, Technical University Munich, Munich, Germany) for cell sorting. This study was Funded by the German Research Foundation (DFG: OO 8/16, OO 8/18, and FOR 2033/2 project B3) and the Translational Medicine Program of the TUM Medical Graduate School.

## Author contributions

RH, MH, AJA, MK, SRM, FH, RI, and RAJO designed experiments. RH, MH, AJA, MK, SRM, and FH acquired, and analyzed data. RH, CS, RI, and RAJO interpreted data. RH, CS, RI and RAJO drafted the manuscript

## Notes

### Competing Interest Statement

The authors have declared no competing interest.

### Summary of Updates

Submission to different journal

