## Supplementary material for "Cathepsin K maintains the number of lymphocytes *in vivo*": Hausinger CTSK Supplemental Files

**Summary**

Flow cytometry dot plots

Figure S1 (to Figure 3)

Figure S2 (to Figure 4)

Figure S3 (to Figure 5)

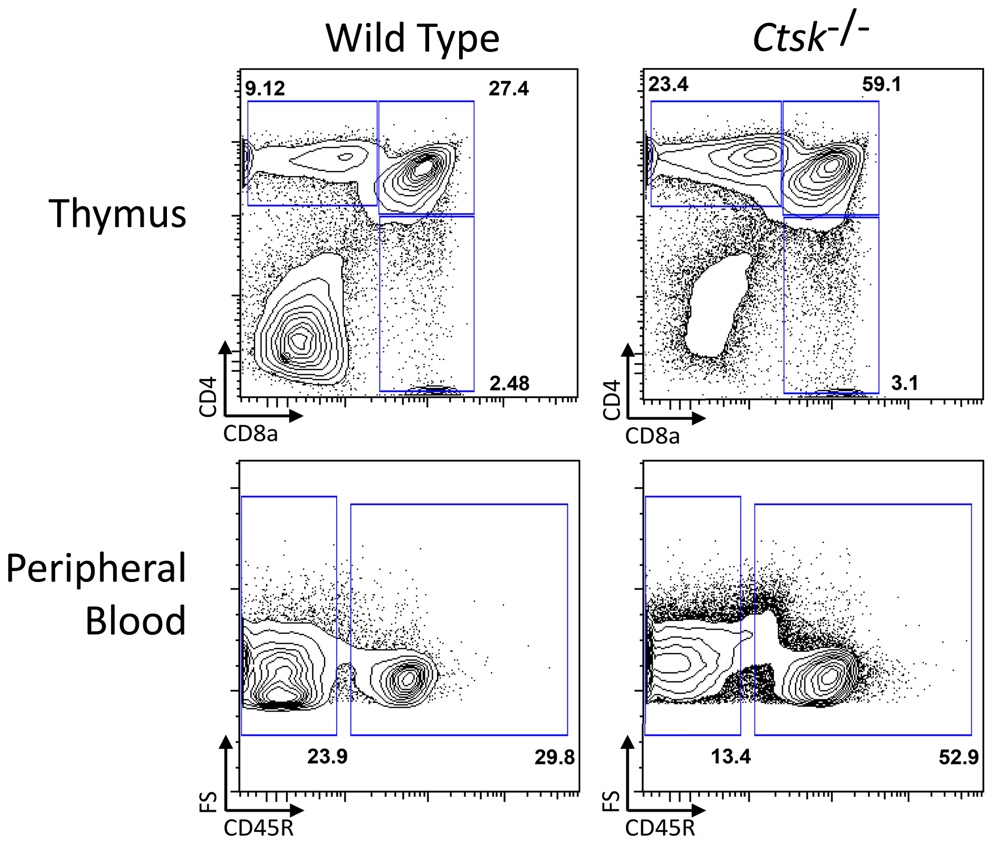

**Figure S1** (to Figure 3). Representative flow cytometry dot plots of the thymus and peripheral blood of 8-12-week old *Ctsk*^-/-^ mice under steady state conditions. Shown is cell staining for CD4 and CD8a in the thymus (top panels) and B cell (CD45R; B220) staining for the peripheral blood (bottom panels). Wild type samples are shown on the left and *Ctsk*^-/-^ samples on the right hand sides.

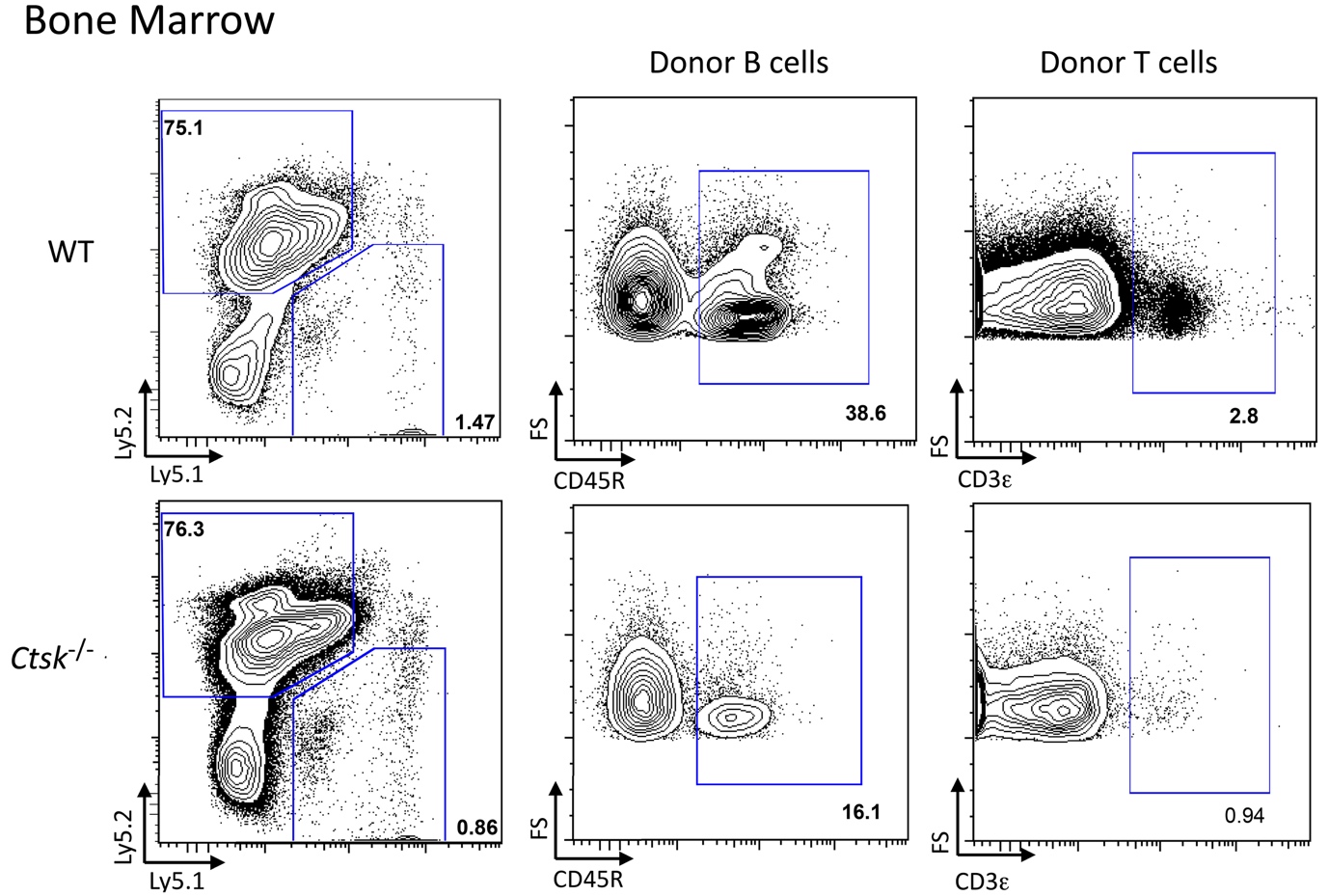

**Figure S2** (to Figure 4). Representative flow cytometry dot plots of the bone marrow 16 weeks after transplantation of wild-type CD45.1 donor cells into CD45.2 wild-type or *Ctsk*^-/-^ recipients. Shown is donor cell staining (CD45.1^-^ CD45.2^+^, left panels), B cell staining (CD45R/B220), middle panels) and T cell staining (CD3ε, right panels). Fluorescence staining was detected using a Beckman-Coulter Cyan, and the analyses were performed using FlowJo software.

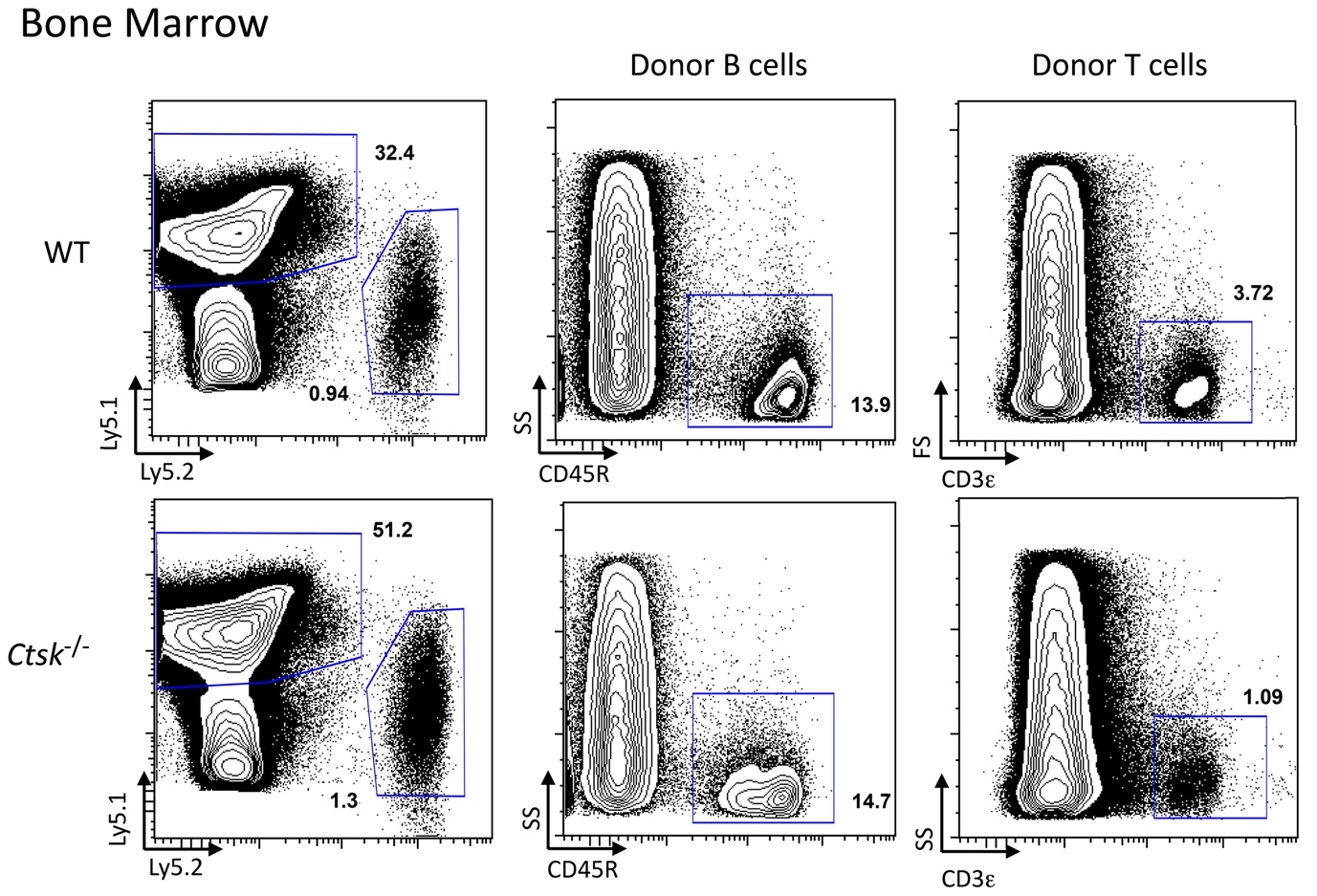

**Figure S3** (to Figure 5). Representative flow cytometry dot plots of the bone marrow 16 weeks after transplantation of CD45.2 WT or *Ctsk*^-/-^ donor cells into wild-type CD45.1 recipients. Shown is donor cell staining (CD45.1^+^ CD45.2^-^, left panels), the B cells (CD45R^+^, middle panels), and T cells (CD3ε^+^, right panels) in WT (upper row) or *Ctsk*^-/-^ (lower row) donors. Fluorescence staining was detected using a Beckman-Coulter Cyan, and the analyses were performed using FlowJo software.
